# *WAPO-A1* is the causal gene of the 7AL QTL for spikelet number per spike in wheat

**DOI:** 10.1101/2021.07.29.454276

**Authors:** Saarah Kuzay, Huiqiong Lin, Chengxia Li, Shisheng Chen, Daniel Woods, Junli Zhang, Jorge Dubcovsky

## Abstract

Improving our understanding of the genes regulating grain yield can contribute to the development of more productive wheat varieties. Previously, a highly significant QTL affecting spikelet number per spike (SNS), grain number per spike (GNS) and grain yield was detected on chromosome arm 7AL in multiple genome-wide association studies. Using a high-resolution genetic map, we established that the A-genome homeolog of *WHEAT ORTHOLOG OF APO1* (*WAPO-A1*) was a leading candidate gene for this QTL. Using mutants and transgenic plants, we demonstrate in this study that *WAPO-A1* is the causal gene underpinning this QTL. Loss-of-function mutants *wapo-A1* and *wapo-B1* showed reduced SNS in tetraploid wheat, and the effect was exacerbated in *wapo1* combining both mutations. By contrast, spikes of transgenic wheat plants carrying extra copies of *WAPO-A1* driven by its native promoter had higher SNS, a more compact spike apical region and a smaller terminal spikelet than the wild type. Taken together, these results indicate that *WAPO1* affects SNS by regulating the timing of terminal spikelet formation. Both transgenic and *wapo1* mutant plants showed a wide range of floral abnormalities, indicating additional roles of *WAPO1* on wheat floral development. Previously, we found three widespread haplotypes in the QTL region (H1, H2 and H3), each associated with particular *WAPO-A1* alleles. Integrating results from this study and previous findings, we show that the *WAPO-A1* allele in the H1 haplotype (115-bp deletion in the promoter) is expressed at significantly lower levels in the developing spikes than the alleles in the H2 and H3 haplotypes, resulting in reduced SNS. Field experiments also showed that the H2 haplotype is associated with the strongest effects in increasing SNS and GNS (H2>H3>H1). The H2 haplotype is already present in most modern common wheats, so it might be particularly useful in durum wheat where H2 is rare.

**AUTHOR SUMMARY:** A region on wheat chromosome 7A has been previously shown to affect the number of spikelets and grains per spike as well as total grain yield in multiple breeding programs. In this study, we show that loss-of-function mutations in the *WAPO1* gene located within this region reduce the number of spikelets per spike and that additional transgenic copies of this gene increase this number. These results demonstrate that *WAPO1* is the gene responsible for the differences in grain number and yield associated with the 7A chromosome region. Among the three main variants identified for this gene, we demonstrate in field experiments that the H2 variant is associated with the largest increases in number of spikelets and grains per spike. The H2 *WAPO1* variant is frequent in bread wheat breeding programs but is almost absent in modern pasta wheat varieties. Therefore, the introgression of the H2 represents a promising opportunity to improve grain yield in pasta wheat breeding programs.

## INTRODUCTION

Wheat is an essential staple crop for global food security. It is highly adapted to a wide variety of climates and production systems, and provides more than 20 % of the calories and protein consumed by the human population [1]. Although further increases in grain yield are required to feed a continuously growing population, historical yield trend studies have shown a decrease in the relative rates of grain yield gains in some wheat growing regions [2]. This has prompted new efforts to understand and improve the productivity of both common (*Triticum aestivum*, genomes AABBDD) and durum wheat (*T. turgidum* ssp. *durum*, genomes AABB).

Identifying genes controlling total grain yield is challenging due to the low heritability of this trait [3, 4]. However, grain yield can be dissected into more discrete yield components with higher heritability. Total grain yield can be partitioned into the number of spikes per unit of area, spikelet number per spike (SNS), grain number per spikelet, and average grain weight. Among these traits, SNS usually exhibits high heritability because it is established early in the reproductive phase when the terminal spikelet is formed [5], limiting the effect of environmental conditions after this point.

A highly significant and stable QTL for SNS was identified on chromosome arm 7AL in multiple genome-wide association studies (GWAS) including a panel of soft red winter wheats in the US [6], panels of European winter wheats [7-9], a panel of US and CIMMYT photoperiod-insensitive spring wheats, and six biparental populations which comprised different wheat market classes [10]. In our previous study [10], we generated two high-resolution genetic maps to delimit this SNS QTL to an 87-kb region (674,019,191 – 674,106,327 bp, RefSeq v1.0) containing four candidate genes. Among these genes, we identified *TraesCS7A02G481600* as the most promising candidate gene, based on the presence of a non-synonymous polymorphism that co-segregated with SNS in biparental populations segregating for different haplotypes in the candidate region [10].

The wheat gene *TraesCS7A02G481600* is orthologous to the *Oryza sativa* (rice) gene *ABERRANT PANICLE ORGANIZATION1* (*APO1*), hence it was designated as *WHEAT ORTHOLOG of APO1* (*WAPO1*). Mutants in the rice *APO1* gene affect panicle branching and spikelet number [11], supporting *WAPO-A1* as a promising candidate gene for the SNS QTL [10]. The rice *APO1* gene and its homolog in *Arabidopsis thaliana* (Arabidopsis), *UNUSUAL FLORAL ORGANS* (*UFO*), encode an F-box protein that is a component of an SCF (**S**kp1– **C**ullin–**F**-box-protein) ubiquitin ligase. This component is important to maintain the activity of LEAFY (LFY), a transcription factor that plays key roles in flowering and floral development [12].

In rice, mutations in *APO1* or *LFY* (also known as *APO2* and *RFL* in rice) result in reductions in the number of branches and spikelets per panicle. The effect is similar in the *apo1 lfy* double mutant suggesting that these two genes act cooperatively to control this trait [13]. Mutations in these genes are also associated with floral abnormalities, but in this case the *apo1 lfy* double mutant shows more severe phenotypes than either of the two single mutants, suggesting that these genes may also play independent roles in floral development [13]. Floral defects in the rice *apo1* and Arabidopsis *ufo* mutants are concentrated in the internal whorls [14, 15].

Rapid changes in *WAPO-A1* allele frequencies during wheat domestication and breeding suggest that this region is relevant to wheat improvement. Three major haplotypes were identified in the 87-kb candidate region – H1, H2, and H3 – each of which associated with different *WAPO-A1* alleles. Haplotype H3 includes the ancestral alleles *Wapo1-A1c* and *Wapo1-A1d*, which differ from each other by two synonymous substitutions, two SNPs in the single intron and one in the promoter, which likely have limited effect on gene function [10]. The H3 haplotype is present in the diploid donor of the A genome (*T. urartu*), cultivated emmer (*T. turgidum* subsp. *dicoccon*) and wild emmer (*T. turgidum* subsp. *dicoccoides*), and at low frequency in modern durum and common wheat varieties. Haplotype H1, present in over 99 % of modern durum wheat lines, has the *Wapo-A1a* allele that is characterized by a 115-bp deletion in the promoter and a change from aspartic acid to asparagine at position 384 (D384N) relative to H3. Haplotype H2, the most frequent in modern common wheat varieties, carries the *Wapo-A1b* allele and differs from the ancestral haplotype by a cysteine to a phenylalanine polymorphism at amino acid position 47 (henceforth, C47F), in a conserved region of the F-box motif. Linkage analysis in six different biparental populations established that the H2 haplotype was associated with higher SNS than both the H1 and H3 haplotypes [10].

Our previous study established *WAPO-A1* as the best candidate gene for the 7AL SNS QTL [10], but functional validation was missing. In this study we demonstrate that *WAPO1* is the gene underpinning the SNS QTL by characterizing loss-of-function mutants and transgenic plants. We also describe the flower abnormalities observed in plants with complete loss of *WAPO1* activity and in transgenic plants with additional *WAPO1* genes. Finally, we characterize the effect of different natural *WAPO-A1* alleles on the number of spikelets and grains per spike and discuss their potential applications in common and durum wheat breeding programs.

## RESULTS

### Loss-of-function EMS and CRISPR mutations in *WAPO1* reduce spikelet number per spike

The EMS induced mutation W216* identified in Kronos mutant line K4222 results in a premature stop codon and the truncation of 51 % of the WAPO-A1 protein. The truncated region includes sequences highly conserved among grasses, suggesting that the truncated protein is either non-functional or has very reduced activity. The modified activity of W216* truncated gene was confirmed by the significant SNS reduction in lines carrying the *wapo-A1* mutant allele relative to those carrying the wild type (WT) allele in three different experiments (Fig 1). In the greenhouse experiment, the average SNS of F_3_ lines homozygous for *wapo-A1* EMS mutant was 1.7 spikelets lower (14 % reduction, *P* < 0.001) than the sister lines homozygous for the WT allele (Fig 1A). A similar reduction of 1.4 spikelets (7.3 % reduction, *P* < 0.001) was observed in a field experiment using homozygous F_4_ sister lines for the *wapo-A1* and WT alleles (Fig 1B).

**Figure 1.**
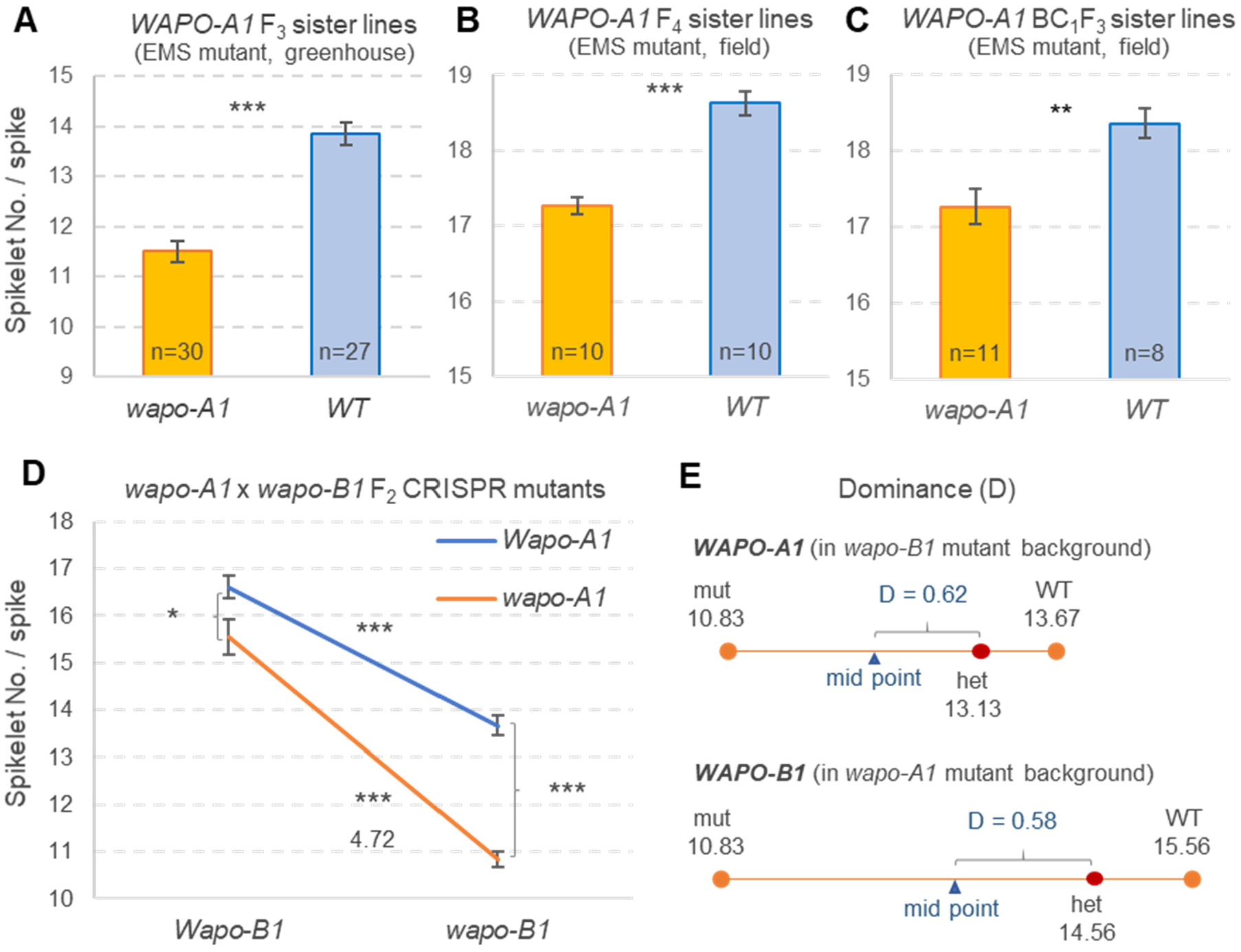
Differences in spikelet number per spike (SNS) between homozygous Kronos sister lines segregating for the EMS induced *wapo-A1* W216* mutation or for CRISPR induced *wapo-A1* and *wapo-B1* mutations. **A**) F_3_ sister lines evaluated in the greenhouse. (**B**) F_4_ sister lines evaluated in the field. (**C**) BC_1_F_3_ sister lines evaluated in the field. (**D**) Interaction graph showing the effect of loss-of-function CRISPR mutants in *wapo-A1* and *wapo-B1* in tetraploid wheat Kronos plants without the transgene. The factorial ANOVA is presented in S2A Table. (**E**) Degree of dominance [16] for *WAPO-A1* (in *wapo-B1* background) and *WAPO-B1* (in *wapo-A1* background). Bars are average SNS and error bars are s.e.m. * = *P* < 0.05, ** = *P* < 0.01 and *** = *P* < 0.001. **= *P* < 0.01, *** *P* < 0.001.

To reduce potential variability originating from background mutations and more accurately determine the effect of the W216* truncation, we crossed the K4222 mutant to WT Kronos twice and selected homozygous BC_1_F_2_ sister lines for an additional field evaluation using their BC_1_F_3_ grains. The BC_1_F_3_ plants homozygous for the *wapo-A1* had an average of 1.1 fewer spikelets per spike relative to the WT sister line (6.0 % reduction, *P =* 0.0035, Fig 1C). Taken together, these three experiments demonstrate that loss-of-function of the A-genome homeolog of *WAPO1* is sufficient to significantly reduce SNS.

To test the effect of the *WAPO-B1* homeolog and validate the results for the EMS induced *wapo-A1* mutant, we generated three independent T_0_ CRISPR-Cas9 transgenic Kronos plants with a guide RNA targeting both homeologs. We identified one line with a “T” insertion at position 510 from the ATG (4-bp upstream of the CGG PAM site) in both *WAPO-A1* and *WAPO-B1*, and selected it for further characterization. This frameshift insertion alters 61.4 % of the protein sequence (starting from amino acid 171), likely resulting in loss-of-function of both *WAPO1* homeologs. We genotyped 110 T_1_ plants derived from the selected transgenic event using the *WAPO-A1* and *WAPO-B1* CAPS markers described (S1 Table). The proportion of lines homozygous for the WT *Wapo-A1* (2.7 %) and/or *Wapo-B1* (4.5 %) alleles was significantly lower than the expected 25 %, suggesting continuous CRISPR editing. This hypothesis was validated by the identification of a novel 5-bp frame-shift deletion starting at position 505 in *WAPO-B1* (4-bp upstream the PAM site) in line T_1_-40, which was re-sequenced and crossed to WT Kronos to eliminate the CRISPR-Cas9 transgene.

We created lines homozygous for all four possible combinations of the *WAPO1* homeologs – WT, *wapo-A1, wapo-B1*, and *wapo1* double mutant – by selecting F_2_ progeny derived from an F_1_ plant without the CRISPR-Cas9 transgene. In a growth chamber experiment, we evaluated the effects of the two homeologs and their interactions on SNS (Fig 1D), heading time and leaf number (S2 Table). The factorial ANOVAs showed highly significant differences for SNS (S2A Table), but non-significant differences for heading date (S2B Table) or leaf number (S2C Table).

Both *wapo-A1* and *wapo-B1* mutants had significantly lower SNS than the WT (*P <* 0.0001, Fig 1D, S2A Table), with larger reductions in *wapo-B1* (3.7 spikelets) than in *wapo-A1* (1.3 spikelets). The transcript levels of the two homeologs were also different, with higher levels of *WAPO-B1* than *WAPO-A1* at different stages of spike development (S1 Figure). The effect of the *wapo-A1* allele on SNS was highest when the B-genome homeolog was homozygous for *wapo-B1*, and vice versa resulting in a highly significant interaction (*P* = 0.0096, Fig 1D, S2A Table). Partial dominance at the *WAPO-A1* (D = 0.61) and *WAPO-B1* (D = 0.58) loci (Fig 1E) indicated partial functional redundancy between the two copies within each of the *WAPO1* homeologs.

### Transgenic plants with additional *WAPO1* copies show delayed heading time and higher SNS

We cloned the genomic regions of *WAPO-A1* from diploid wheat *T. monococcum* (TmDV92) and tetraploid Langdon with the natural (LDN-C47) and edited allele (LDN-F47) and transformed them into Kronos to test their effects on heading date and SNS. The cloned genomic regions include the intron and regulatory regions (see Materials and Methods), and their coding regions carry different non-synonymous polymorphisms (Table 1).

**Table 1.**
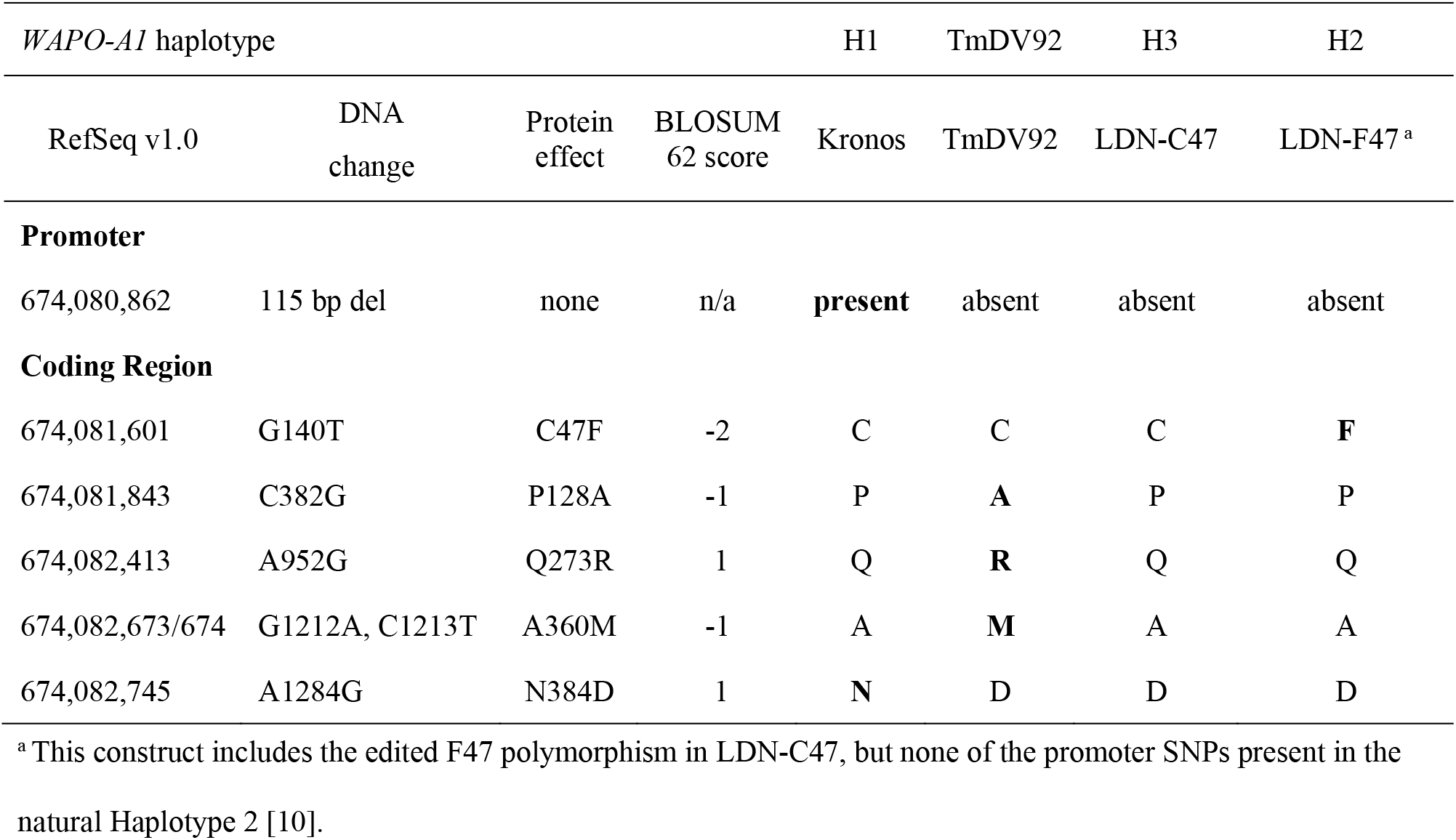
Comparison of regulatory and coding *WAPO-A1* regions in wild type Kronos (H1 haplotype) and the three genomic constructs used in the transgenic plants.

We obtained four independent transgenic events for TmDV92, three for LDN-C47, and five for the LDN-F47 derived allele, and evaluated their corresponding T_1_ progeny in a greenhouse experiment (Fig 2). Transgenic plants for all three constructs showed similar average increases in SNS relative to the WT: LDN-C47 (13.4 %), TmDV92 (15.1 %) and LDN-F47 (15.5 %) (Fig 2). Increases in SNS were significant for two TmDV92 events, one LDN-C47 event (Fig 2A) and four LDN-F47 independent events relative to the WT control (Fig 2C).

**Figure 2.**
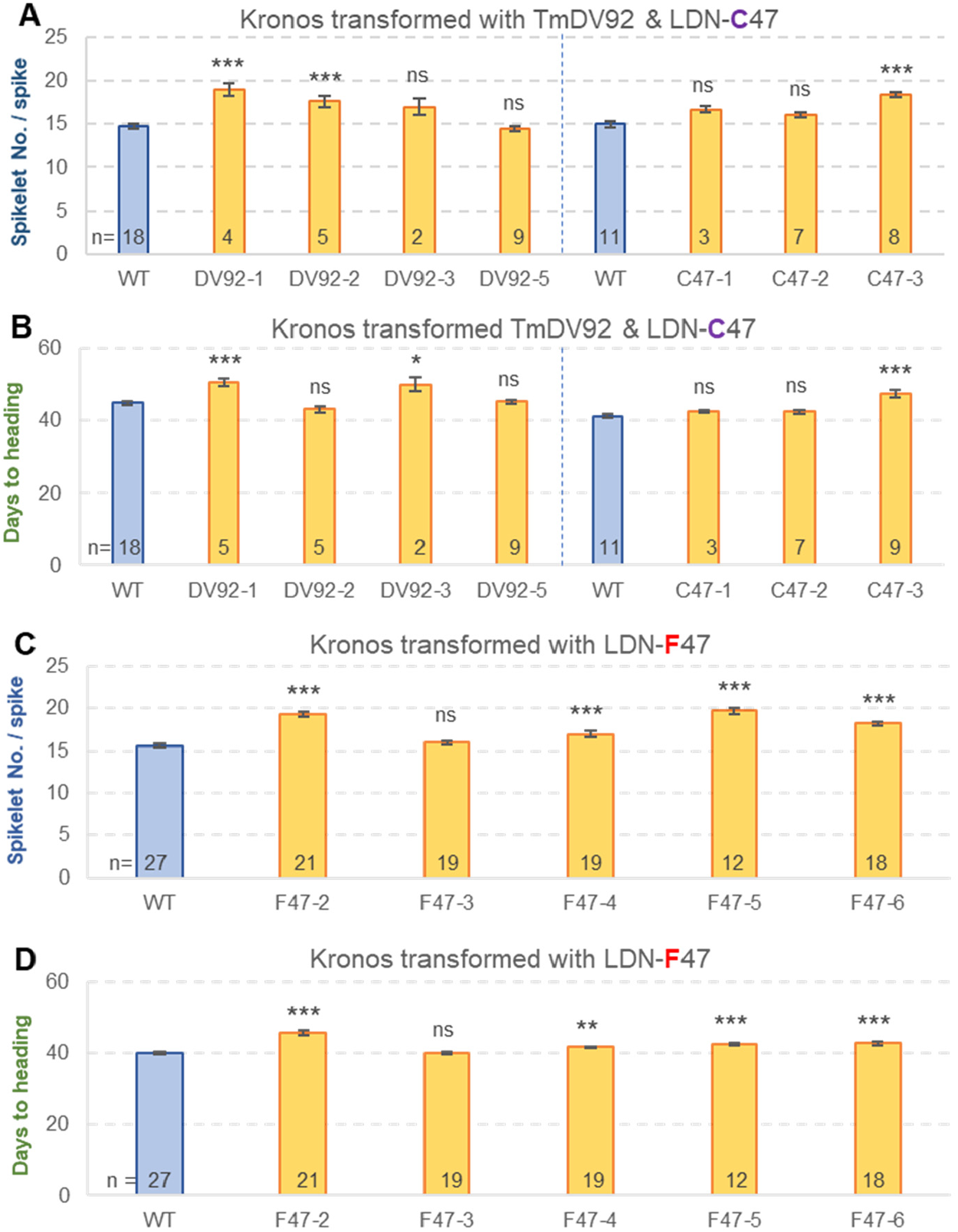
Effect of the *WAPO-A1* alleles from TmDV92 and tetraploid LDN on spikelet number per spike and heading date. (**A-B**) Greenhouse experiment comparing four independent TmDV92 transgenic lines and three independent lines carrying the LDN-C47 allele with their respective non-transgenic Kronos sister lines (pooled from the different events). (**C-D**) Separate greenhouse experiment comparing five independent transgenic lines carrying the LDN-F47 allele with the pool of non-transgenic sister lines. (A and C) Spikelet number per spike. (B and D) Heading date. Bars are averages and error bars are s.e.m. Numbers inside bars indicate number of plants and asterisks indicate *P* values (Dunnett test) relative to the pool of non-transgenic sister lines carrying the same construct. * = *P* < 0.05, ** = *P* < 0.01 and *** = *P* < 0.001.

The effects of the transgenes on heading date were smaller and slightly less significant than those for SNS. For heading date, a 7.0 % delay was observed in transgenic plants with the LDN-C47 construct, a 5.4 % delay for TmDV92, and a 6.3 % delay for LDN-F47 (Fig 2B and D). Heading date was positively correlated with SNS among the T_1_ lines segregating for each of the three constructs (*R =* 0.49-089, *P* <0.01). For event F47-2, which showed the strongest delay in heading time and high SNS (Fig 2C-D), we confirmed the presence of significantly higher transcript levels of *WAPO-A1* in the developing spikes of the transgenic plants relative to a WT sister line (S2 Figure). Taken together, these results demonstrate that the presence of additional copies of *WAPO1* in the transgenic plants increases SNS and delays heading date.

We also evaluated SNS and heading date in transgenic plants expressing the *Wapo-A1c* (C47) and *Wapo-A1b* (F47) coding regions, each fused to a C-terminal MYC tag and driven by the constitutive maize *UBIQUITIN* promoter. Using qRT-PCR, we measured *WAPO-A1* transcript levels in the leaves of 11 independent events for UBI::C47:MYC and 14 events for UBI::F47:MYC (S3A Figure). Transgenic lines for all events showed higher *WAPO-A1* transcript levels in the leaves than non-transgenic Kronos, where expression of *WAPO-A1* was not detected. We selected events with the highest transcript levels for further evaluation; E35 for UBI::C47:MYC (11.5-fold *ACTIN*) and E53 for UBI::F47:MYC (12.3-fold *ACTIN*). The T_2_ transgenic progeny of these lines headed significantly later than the non-transgenic sister lines (S3B Figure) but showed no significant differences in SNS (S3C Figure).

### CRISPR *wapo1* mutant shows altered floral morphology

In addition to producing fewer SNS, *wapo1* double mutant displayed a great diversity of floral abnormalities and produced a limited number of grains. To establish the frequency of the different floral abnormalities, we dissected 91 florets from 32 spikelets located at different positions along the spike from 14 different *wapo1* mutant plants. While glumes, lemmas and paleas were normal, multiple abnormalities were observed in other floral organs. The lodicules, which are normally two per floret, varied from zero to four (average 1.95, Fig 3A) and were frequently fused with stamens (19.1 %), leaf-like tissue (22.5 %) or both (5.6 %, Fig 3B-D). The number of stamens per floret was lower than three in 42 % of the florets (Fig 3 A-D), resulting in an average of 2.15 stamens per floret. Stamens were frequently fused with each other or with ovaries (6.6 %), lodicules (18.9 %), leaf-like tissue (9.2 %) or combinations of two of the previous three categories (4.6 %, Fig 3B and C). Most florets had one pistil with one ovary, but 43 % of the florets had multiple pistils, likely due to loss of floret meristem determinacy. In 28.4 % of the florets, the ovaries were fused with leafy tissue (Fig 3D).

**Figure 3.**
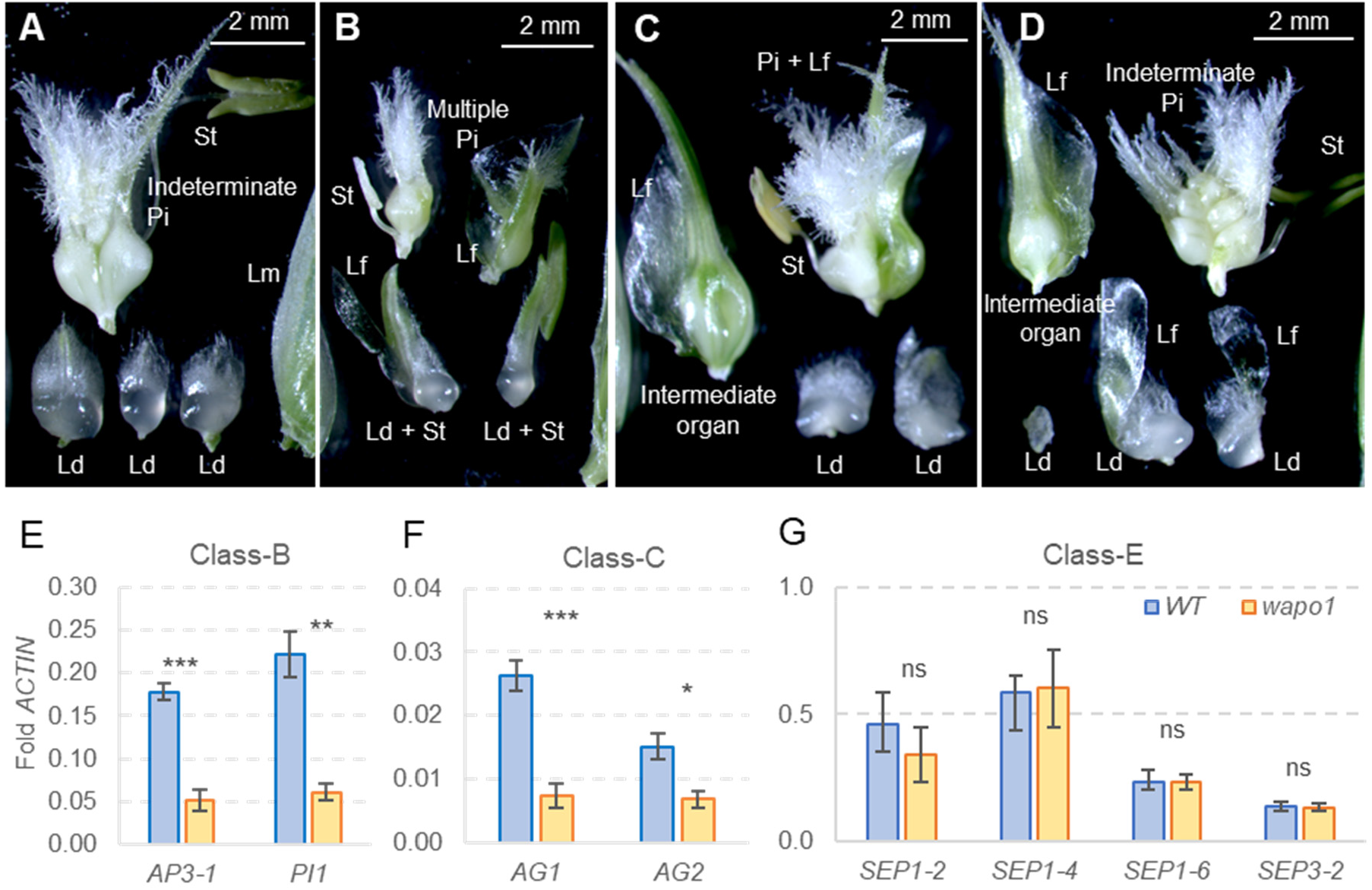
Floral abnormalities and expression profiles of Kronos *wapo1* loss-of-function CRISPR mutant (*wapo-A1 wapo-B1*). (**A**) Mutant T_1_-101, floret one from basal spikelet showing three lodicules, one stamen and two fused pistils. (**B**) Mutant T_1_-101, 2^nd^ floret from 3^rd^ most distal spikelet. Lodicules fused with stamens and leafy tissue. Indeterminate pistils fused with leafy tissue. (**C**) Mutant T_1_-13, 2^nd^ floret of basal spikelet. One lodicule fused with leafy tissue, and an intermediate organ fused with leafy tissue, and pistil fused with leafy tissue. (**D**) Mutant T_1_-13, 3^rd^ floret of basal spikelet. Lodicules with elongated leafy tissue, intermediate organ and indeterminate pistils. Lm = lemma, Pa = palea, Ld = lodicule, St = stamen, Pi = pistil, Lf = fused leafy tissue, + = fused organs. (**E-G**) Expression analysis of class-A, -B, -C and -E MADS-box genes in Kronos and *wapo1* mutants (MADS-box gene nomenclature is based on [17]). Primers are listed in S1 Table. Bars represent averages of four replicates and error bars are s.e.m. Each replicate is a pool of 12 meristems at the stamen primordia stage (∼W4.0 in Waddington scale). ns = not significant, * = *P* < 0.05, ** = *P* < 0.01, and *** = *P* < 0.001.

Since most of the floral abnormalities in *wapo1* were detected in the internal whorls (lodicules, stamens and pistils), we explored the effect of the *wapo1* mutant on the expression of class-B, -C an -E MADS-box floral genes in developing spikes at the stamen primordia stage. Relative to the WT, the *wapo1* mutant showed a significant down regulation of class-B (*AP3-1* and *PI1*) and class-C (*AG1* and *AG2*) MADS-box genes (Fig 3E-G), but no significant differences were detected for the control *SEPALLATA* MADS-box genes *SEP1-*2, *SEP1-*2, *SEP1-4, SEP1-6*, or *SEP3-1* (Fig 3H).

### *WAPO1* transgenic plants exhibit abnormal spike and floral phenotypes

Among T_1_ plants segregating for the three transgenic *WAPO1* copies driven by their native promoters, the events showing significant differences in SNS also displayed spike abnormalities (Fig 4). One unusual phenotype was the presence of naked pistils at the base of spikelets located at basal positions of the spike (Fig 4A-C). This abnormality was observed in some plants from events TmDV92-1, LDN-C47-3, LDN-F47-2 and LDN-F47-6 (Fig 4A-C).

**Figure 4.**
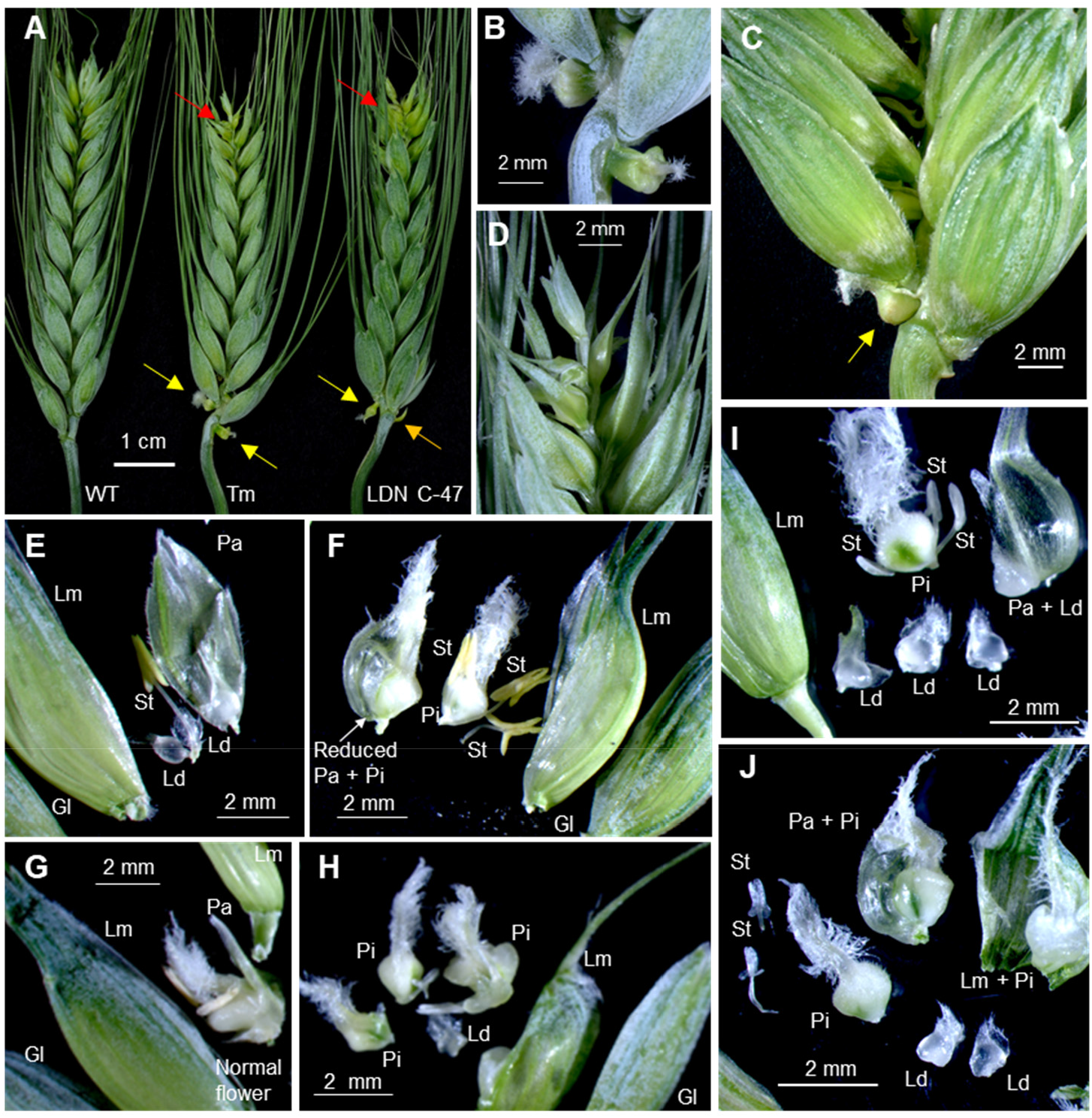
Floral abnormalities in Kronos plants transformed with genomic regions of *WAPO-A1*. (**A**) Spikes of non-transgenic control (WT) and transgenic lines TmDV92-1 and LDN-C47-3. (**B**) Detail of the naked pistils below the basal spikelets in TmDV92-1 (yellow arrows in A). (**C**) Naked pistil below 2^nd^ spikelet in LDN-F47. (**D**) Detail of the abnormal distal end of the spike of TmDV92-1 (red arrows in A). (**E** and **F**) LDN-C47-3. (**E**) First floret in the 5^th^ spikelet from the base: reduced palea, one stamen and no pistil. (**F**) Second floret in 11^th^ spikelet from the base: no lodicules and extra pistils fused with small palea. (**G** and **H**) TmDV92-1. (**G**) First floret from basal spikelet: small palea and normal flower. (**H**) 17^th^ spikelet from the base: one lodicule, two stamens and indeterminate pistils. (**I** and **J**) LDN-F47-2. (**I**) Third floret from basal spikelet: 4 lodicules one fused to palea. (**J**) Spikelet 19^th^: 2 small stamens and three pistils (one fused to the palea and one to the lemma. Lm = lemma, Pa = palea, Ld = lodicule, St = stamen, Pi = pistil.

A more frequent spike abnormality was the presence of smaller, densely packed spikelets in the distal region of the spike, ending in a small terminal spikelet. For simplicity, this spike abnormality is referred to hereafter as small terminal spikelet (Fig 4A red arrows and D). For event LDN-F47-3, which showed no significant differences in SNS, none of the T_1_ plants had small terminal spikelets, whereas this phenotype was observed in 91.6 % of the T_1_ plants of LDN-F47-5, the event with the largest increase in SNS (26 %). The other three LDN-47 events, showed significant but smaller increases in SNS (9 to 24 %) than the WT sister lines, and had small terminal spikelets in approximately half of the T_1_ lines (47 to 50 %). A strong correlation was detected between the proportion of spikes showing small terminal spikelets and the percentage increase in SNS (*R =* 0.85, n = 5) suggesting that the increases in SNS in these transgenic plants were associated with the additional smaller spikelets in the distal region of the spike.

Dissection of spikelets at different positions of the spike revealed increasing floral abnormalities from the basal positions to the apical region in LDN-C47-3 (Fig 4 E-F), TmDV92-1 (Fig 4 G-H), LDN-F47-2 (Fig 4 I-J), and LDN-F47-4 (S4 Figure). The abnormal floral structures included (i) small paleas, (ii) variable numbers of lodicules, stamens and pistils per floret, and (iii) frequent fusions among these three organs and with lemmas and paleas (Fig 4 E-J). Transgenic line LDN-F47-4 had the most extreme phenotype showing multiple naked pistils, which were larger at the basal spikelets and transitioned to bracts in the more distal spikelets (S4A-B Figure). Spikelets near the terminal spikelet had higher numbers of pistils and reduced numbers of other organs (S4C Figure). By contrast, basal spikelets of LDN-F47-4 showed larger glumes, lemmas and stamens, particularly in the distal florets (S4D-G Figure).

We also characterized the spikes and florets of transgenic plants transformed with UBI::C47:MYC (E35) and UBI::F47:MYC (E53) constructs. Unexpectedly, these transgenic plants showed milder spike and floral defects (S5 Figure) than the plants transformed with *WAPO-A1* genomic regions (Fig 4 and S4 Figure), possibly due to the fused MYC-tag. Most UBI::C47:MYC and UBI::F47:MYC plants showed small terminal spikelets (S5A-B Figure) and a small proportion of plants had naked pistils on the underside of the basal spikelets (S5C Figure). While most flowers of the UBI::C47:MYC and UBI::F47:MYC plants showed small and sometimes curved paleas, organ numbers were similar to the WT. The most frequently observed organ fusions were between paleas and lodicules, while fusions between stamens were (S5D-E Figure). No obvious abnormalities were detected in pistils.

We did not observe significant phenotypic differences between the transgenic plants transformed with the UBI::C47:MYC (E35) and UBI::F47:MYC (E53) or among plants transformed with the three different genomic constructs, suggesting that all these alleles encode active proteins. The large variability among the different events for the same construct precluded the detection of smaller quantitative differences among constructs.

### Natural variation in *WAPO-A1* is associated with changes in SNS

The characterization of *WAPO-A1* natural variants complemented the information obtained from the induced mutations and transgenic events described above. In our previous study, we established that the *WAPO-A1* allele in the H2 haplotype was associated with higher SNS than the H1 and H3 haplotypes [10], but we did not determine the relative effects of the H1 and H3 haplotypes on SNS. To address this question, we performed two field experiments organized in a split plot-RCBD design in 2020 (Fig 5A, S3A Table) and 2021 (Fig 5B-D, S3B-F Tables). Both experiments had 10 replications, each including eight families as main plots and homozygous H1 and H3 sister lines as subplots. In the 2020 experiment, average SNS for plants homozygous for the H3 haplotype was 6 % higher than for those homozygous for the H1 haplotype in 2020 (1.3 spikelets) and 7.3 % higher in 2021 (1.6 spikelets). The overall differences in SNS were highly significant both years (*P* < 0.0001, Fig 5A and B, S3A and B Tables). Within individual families, differences in SNS between H1 and H3 were significant for six out of the 8 families in 2020 and for all families in 2021 (S3A and B Tables).

**Figure 5.**
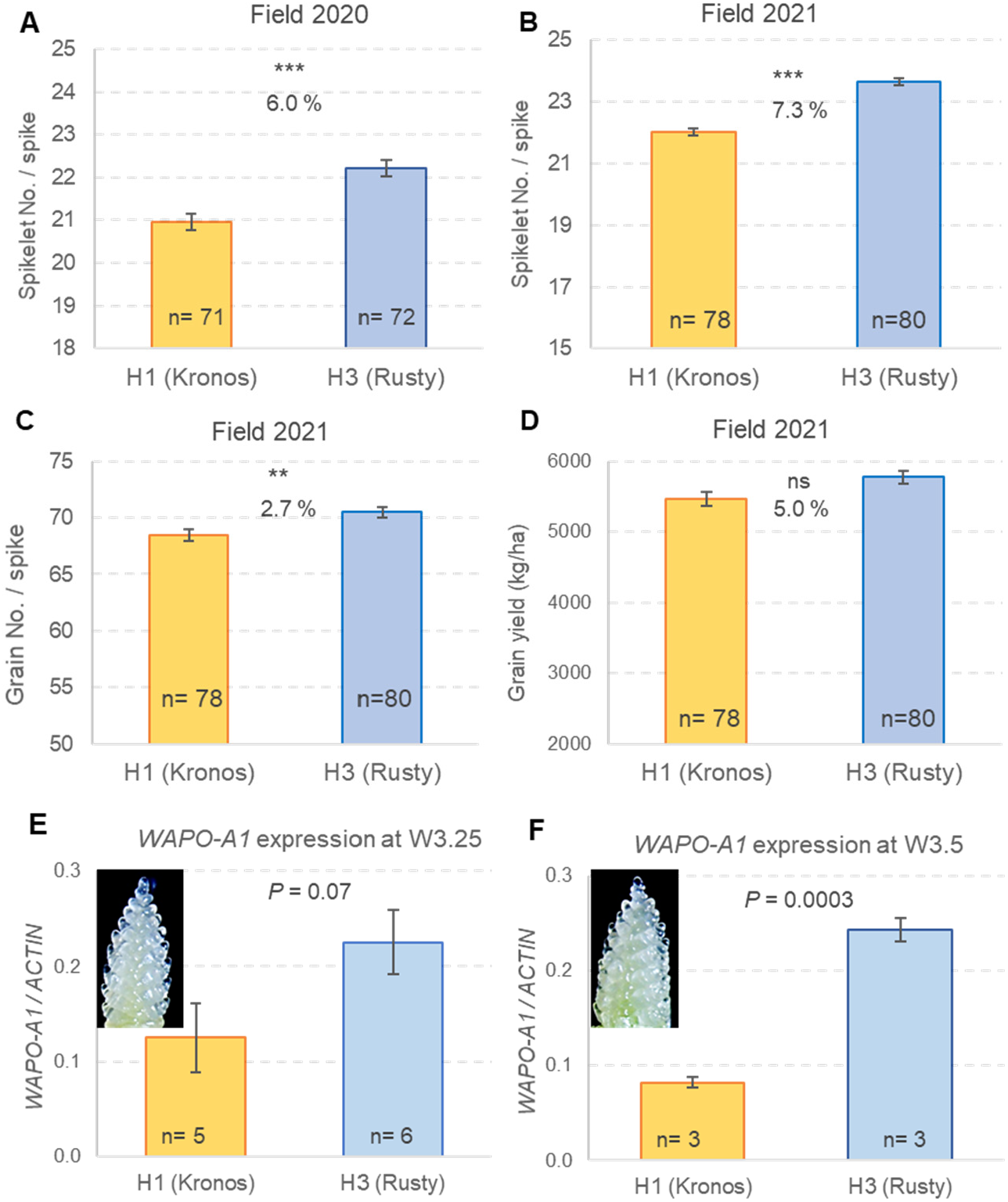
Effect of *WAPO-A1* haplotypes H1 (Kronos) and H3 (Rusty) on SNS and *WAPO-A1* transcript levels. (**A**) 2020 Split-plot RCBD field experiment. (**B-D**) 2021 Split-plot RCBD field experiment. (**B**) Spikelet number per spike. (**C**) Grain number per spike. (**D**) Grain yield (kg/ha) (**E - F**) Two independent qRT-PCR experiments comparing H1 and H3 transcript levels of *WAPO-A1* relative to *ACTIN* using the 2^▵Ct^ method. **C**) Pools of developing spikes collected when lemma primordia were present (W3.25). (**D**) Pools of developing spikes collected when floret primordia were present (W3.5). Bars represent averages and error bars are s.e.m. Numbers inside the bars indicate the number of replications. ns = not significant, * = *P* < 0.05 and *** = *P* < 0.001. A combined ANOVA for C and D using experiment as block showed significant differences in expression between H1 and H3 (*P* = 0.0128).

In the 2021 experiment, we had more grains available so we were able to use small plots (1.86 m^2^) as experimental units to provide a preliminary estimate of grain yield. In this experiment, the H3 genotype showed a 2.7 % increase in GNS (*P =* 0.0085, Fig 5C, S3C Table) and a 1.7 % decrease in kernel weight (*P =* 0.0319, S3D Table) relative to H1. The balance of this negative correlation was a 5.0 % increase in grain yield, but the difference was marginally not significant (*P* = 0.0525, Fig 5D, S3E Table). In this experiment, we did not detect significant differences in heading time (*P =* 0.5987, S3F Table).

To test if the differences in SNS between H1 and H3 haplotypes were associated with differences in the expression levels of *WAPO-A1*, we performed two independent qRT-PCR experiments comparing homozygous H1 and H3 sister lines derived from HIF #120. Developing spikes from the main tiller were collected at the lemma primordia stage (W3.25) for the first experiment and at the floret primordia stage (W3.5) for the second experiment (Fig 5E and F). In the first experiment, transcript levels in H3 were 80 % higher than in H1, but the difference was not significant (*P* = 0.07, Fig 5E). In the second experiment, transcript levels in H3 were 197 % higher than in H1 and the differences were highly significant (*P =* 0.0003, Fig 5F). Since the two experiments showed the same trend, we performed a combined ANOVA using experiment as block. This analysis confirmed the significantly higher *WAPO-A1* transcript levels in H3 than in H1 (*P* = 0.0128).

In the same field used for comparing the H1 and H3 haplotypes in 2021, we also compared the relative effects of the H2 and H1 haplotypes in near isogenic lines (NILs) of tetraploid wheat Kronos and hexaploid wheat GID4314513 (a high-biomass line from CIMMYT). In the tetraploid BC_3_F_3_ NILs, we did not find significant differences in heading date (Fig 6A), but observed 17.6 % more spikelets per spike (*P* < 0.0001, Fig 6B) and 6.7 more grains per spike (*P* = 0.025, Fig 6C) in the H2-NILs than in the H1-NILs (S4 Table). Grain weight per spike (grain number x grain weight) was 5.8 % higher in the H2-NILs than in the H1-NILs, but the differences were not significant (*P* = 0.11, Fig 6D, S4 Table). Total grain yield was not estimated in this experiment because experimental units were single rows.

**Figure 6.**
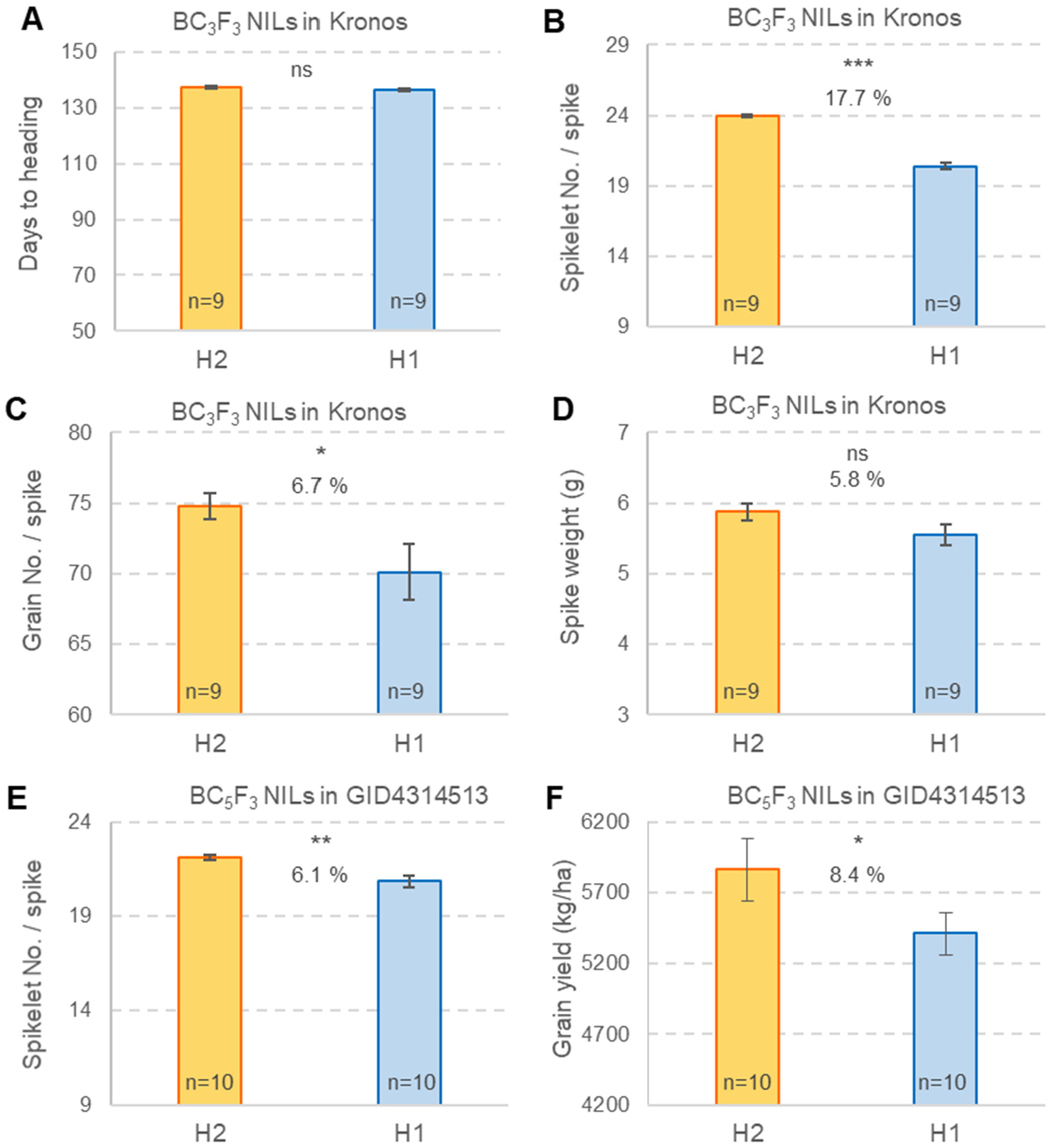
Effect of *WAPO-A1* haplotypes H2 introgressed into tetraploid Kronos (H1) and high-biomass hexaploid line GID4314513 (H1) in field experiments (2021). (**A-D**) Homozygous BC_3_F_3_ sister lines (CRD, n = 9, 1-m rows, 10 spikes measured per row). (**A**) Heading date. (**B**) Spikelet number per spike. (**C**) Grain number per spike. (**D**) Grain weight per spike. (**E-F**) Homozygous BC_5_F_3_ sister lines (RCBD, n = 10, 1.86 m^2^ plots, 4 spikes measured per plot). Error bars are s.e.m. ns = not significant, * = *P* < 0.05, ** = *P* < 0.01, *** = *P* < 0.001.

In the hexaploid BC_5_F_3_ NILs, we detected a 6.1 % increase in SNS in the H2-NILs relative to the H1-NILs (*P* = 0.0092, Fig 6E) and an 8.4 % increase in grain yield that was marginally significant (*P* = 0.0497, Fig 6F, S4E-F Table 4). The H2 haplotype was also associated with non-significant differences in GNS (+2.7 %), number of grains per spikelet (−3.3 %), thousand kernel weight (+1.8 %) and grain weight per spike (+4.4 %, S4G-J Table).

## DISCUSSION

In our previous study, we identified *WAPO-A1* as the leading candidate gene for the 7AL SNS QTL based on two high resolution genetic maps [10]. In this study, we demonstrate that *WAPO1* is both necessary and sufficient to increase SNS in wheat and, therefore, that *WAPO-A1* is the causal gene underpinning the 7AL QTL for SNS.

### Potential mechanisms involved in *WAPO1* effect on SNS

Differences in *WAPO-A1* transcript levels in different genotypes correlate well with differences in SNS. The reduced expression of the *WAPO-A1* allele (115-bp promoter deletion) in H1 relative to the H2 [10] and H3 haplotypes (Fig 5) is associated with lower SNS in H1 relative to the other two haplotypes. Similarly, the higher transcript levels of *WAPO-B1* relative to *WAPO-A1* during spike development is associated with stronger effects of *wapo-B1* mutation on SNS relative to *wapo-A1* (Fig 2B). These results, together with the partial codominant effect of *WAPO1* on SNS (Fig. 1E) and the increased SNS in transgenic plants with additional genomic copies of *WAPO-A1* (Fig 2), support the hypothesis that higher *WAPO1* transcript levels can drive increases in SNS.

Increases in SNS imply a delay in the transition of the inflorescence meristem (IM) into a terminal spikelet. Three indirect lines of evidence suggest that *WAPO1* expression in the wheat IM is likely involved in this mechanism. First, *in situ* hybridization experiments in rice have shown that *APO1*, the rice ortholog of *WAPO1*, is expressed in the IM [11]. Second, we have previously shown by qRT-PCR that *WAPO-A1* is expressed at higher levels in a distal section of early developing spikes than in the more basal regions [10]. Finally, we show that transgenic plants with constitutive expression of *WAPO-A1* driven by the *UBIQUITIN* promoter have more compact spike tips and smaller terminal spikelets (Fig. 4), confirming an effect of this gene on the distal part of the spike.

In rice, APO1 physically interacts with LFY (APO2), and the two genes act cooperatively to control spikelet number per panicle [13]. Similar *in vitro* and *in vivo* interactions have been reported between the Arabidopsis homologs UFO and LFY [12, 18]. Mutants for both genes show similar defects in inflorescence structure and morphology, suggesting that these two genes also act cooperatively to promote floral-meristem identity [19]. Based on the rice and Arabidopsis results, we hypothesize that *LFY* may be involved in the mechanism by which *WAPO1* regulates SNS in wheat. It would be interesting to investigate the interactions between these two genes and MADS-box genes from the *SQUAMOSA* and *SVP* clades, which have been previously shown to also affect SNS in wheat [20, 21].

### Floral and spike abnormalities in mutant and transgenic *WAPO-A1* plants

In addition to its binding to the *AP1* promoter, LFY has been shown to bind to regulatory regions of *APETALA3* (*AP3*, a class-B floral gene) and *AGAMOUS* (*AG*, a class-C floral genes) in Arabidopsis [12, 22-24]. The activation of *AP3* expression requires the activity of both *LFY* and *UFO*, which likely explains the reported downregulation of class-B genes in the *ufo* mutant in Arabidopsis [12, 15] and class-C genes in the *apo1* mutant in rice [14]. In wheat, we also observed downregulation of class-B and -C genes but not class-E genes in the developing spikes of the *wapo1* double mutant relative to the WT, suggesting a conserved molecular mechanism.

In Arabidopsis, class-B and class-C MADS-box genes are associated with the identity of petals, stamens and pistils, which explains the stronger abnormalities detected in the *ufo* mutants in the three inner floral whorls. It has been also shown that ectopic expression of both *PI* and *AP3* rescues the floral organ identity defects of the Arabidopsis *ufo* mutant [25]. Stronger floral defects in the inner whorls of the floret than in glumes, lemmas and paleas has been reported in rice [14] and the wheat *wapo1* mutant in this study. The related floral defects observed in the *ufo, apo1* and *wapo1* mutants suggest functional conservation of these homologous genes across the monocot eudicot divide.

Transgenic wheat plants with additional *WAPO1* genomic copies shared some floral aberrations with the *wapo1* mutant, including differences in the number of lodicules, stamens and pistils and fusion among organs. These results suggest that decreases or increases in *WAPO1* expression can lead to abnormal development in the three inner whorls of the floret. Transgenic *WAPO-A1* plants, both with the native and the *UBI* promoter showed additional phenotypes not observed in the *wapo1* mutant, including reduced and sometimes curved paleas (homologous to sepals), a compact spike tip with a small terminal spikelet, and a significant delay in heading time. We do not know if these additional phenotypes in the transgenic plants are an indirect effect of the elevated *WAPO1* expression levels, or the result of ectopic expression due to the *UBI* promoter or to the absence of important regulatory elements in the cloned native promoter region.

A less frequent but unusual phenotype in the transgenic *WAPO1* plants was the presence of naked pistils, which have been also described in the Arabidopsis *ufo* mutants. In the *ufo* mutants, structures resembling normal pistils were formed at the end of primary and coflorescence shoots [15, 26]. The naked pistils in the transgenic wheat plants were more frequent in basal positions of the spike and appeared adjacent to the base of the spikelets. Paired spikelets or branches are also more frequent in the basal spikelets in other wheat mutants such as *Branched head AP2/ERF* transcription factor [27, 28] and *ppd1* [29]. Based on the terminal position of the naked pistils in Arabidopsis *ufo* mutants, we hypothesize that the naked pistils in the transgenic wheat plants may be located at the end of a short branch including one lateral spikelet.

### Effects of different *WAPO-A1* natural alleles and their potential applications in wheat breeding

Our previous study showed that the H2 haplotype was associated with larger increases in SNS than the H1 and H3 haplotypes in several segregating populations. The favorable effect of H2 on SNS was observed in spring and winter wheats and in different wheat market classes, but these increases not always resulted in increases in grain yield. In genotypes or environments with insufficient resources to fill the extra grains, increases in GNS were offset by decreases in grain weight. However, when H2 was present in well adapted and productive genotypes grown in favorable environments, the increases in SNS were associated with increases in grain yield [10]. In this study, we also observed significant increases in grain yield when the H2 haplotype was introgressed into a highly productive line from the high-biomass program from CIMMYT. Under the favorable irrigation and fertilization conditions used in this study, the increases in SNS associated with the H2 haplotype were associated with an 8.4% increase in total grain yield. This is a promising preliminary result, but needs to be further validated with larger yield trials in different genetic backgrounds.

An increase in the frequency of the H2 haplotype was previously described in common wheat, from less than 50 % in old landraces to more than 80 % in modern wheat varieties [10]. This suggests that selection for high SNS, GNS, or grain yield may have accelerated the increases in H2 frequencies. By contrast, in tetraploid wheat the H1 haplotype replaced the H3 haplotype and became almost fixed in modern durum varieties (99 %) [10]. This increase in H1 frequency cannot be explained by selection for higher SNS or GNS, because our field experiment showed that H1 has significantly lower SNS and GNS than H3 (Fig 5). Since durum wheat has much larger grains than common wheat, we speculate that selection for larger grains may have resulted in an indirect selection for reduced grain number, favoring the H1 allele. This indirect selection was likely driven by the negative correlation between grain number and weight, a trade off hat is magnified in source-limited varieties and/or suboptimal environments.

The low frequency of H2 in durum wheat may be the result of the same indirect selection for fewer and larger grains, but it is also possible that H2 was never properly tested in modern durum wheat varieties. The H2 haplotype was already rare in cultivated emmer, where only one H2 accession (from Syria) was identified in a survey of 364 durum wheat varieties [10]. In this study, we show that the transfer of H2 from common wheat into the tetraploid wheat Kronos was associated with significant increases in SNS and GNS. A similar favorable effect of H2 relative to H1 on SNS was previously reported in a segregating population generated from a cross between cultivated emmer and durum wheat in field trials in Fargo (ND) [10]. These are promising results, but they still require validation in different durum varieties and larger yield trials. To facilitate the introgression of the H2 haplotype in durum wheat, we deposited the Kronos NIL with the H2 introgression in the National Small Grain Collection (PI number pending).

In summary, the validation of *WAPO-A1* as the causal gene of the 7AL SNS QTL and of the positive effects of the H2 haplotype on spikelet and grain number per spike, provides wheat breeders a new tool to improve sink traits without significantly affecting heading time. We hypothesize that the introgression of the H2 haplotype into varieties that are not limited in source traits (e.g. varieties with high biomass) can result in increases in total grain yield in favorable environments.

## MATERIALS AND METHODS

### Ethyl methanesulfonate (EMS) induced mutations in *WAPO-A1*

We screened a database of sequenced EMS induced mutations in the tetraploid wheat variety Kronos [30] using the sequence of *WAPO-A1* (*TraesCS7A02G481600*). For the A-genome homeolog (*WAPO-A1*), we identified line K4222 carrying a mutation that results in the replacement of a tryptophan by a premature stop codon at position 216 of the protein (W216*). For the B-genome homeolog (*WAPO-B1*), we identified five mutations that resulted in amino acid changes but none generated premature stop codons or altered splicing sites; hence, our subsequent studies only included the *WAPO-A1* mutant. We crossed the K4222 mutant to the non-mutagenized Kronos, self-pollinated the F_1_, and from the segregating F_2_ plants we selected sister lines homozygous for the mutant (*wapo-A1*) and the WT alleles (*Wapo-A1*). The derived F_3_ plants were evaluated in the greenhouse and the F_4_ plants in the field. To further reduce background mutations, we generated BC_1_F_2_ homozygous sister lines by backcrossing the F_1_ with Kronos, self-pollinating the BC_1_, and selecting homozygous plants in the next generation. The derived BC_1_F_3_ grains were planted in a field experiment.

For the greenhouse experiment, we used two F_3_ plants per pot (3.8 L) and measured average SNS per plant. In the first field experiment, we planted homozygous mutant and WT F_4_ sister lines in a complete randomized design (CRD) with 10 replicates per genotype. Each replicate included a row of 2-5 plants spaced 0.3 meters apart. Average SNS per row was calculated by measuring SNS from 3 spikes per plant. In the second field experiment, we planted BC_1_F_3_ plants 0.3 m apart segregating for the *WAPO-A1* alleles. Genotyping of these plants revealed 11 homozygous *wapo-A1* and 8 homozygous WT within the same BC_1_ family #54. For each plant, we determined the average SNS from four spikes. Both field experiments were conducted at the University of California Experimental Field Station in Davis, referred to hereafter as UC Davis, during the months of October 2019 and June 2020.

### Generation of *wapo-A1 wapo-B1* double mutants using CRISPR-Cas9

To characterize better the function of *WAPO1*, we edited both homeologs of the tetraploid variety Kronos using CRISPR-Cas9 [31]. We designed one guide RNA between positions 494 and 512 from the starting ATG of the coding region (S1 Table) to induce double-strand breaks in the first exon of both *WAPO-A1* and *WAPO-B1*. This guide RNA was then cloned into a vector which included the Cas9 gene and a *GRF4-GIF1* chimera that increases wheat regeneration efficiency for *Agrobacterium-*mediated transformation [32]. Three independent T_0_ transgenic Kronos plants were obtained from the UCD Plant Transformation facility and were screened for mutations by next generation sequencing (NGS) and restriction enzyme digestion. For the NGS screen, we used primers g641-NGS-F and R that amplify both genomes (S1 Table) and analyzed the data using CRISgo (https://github.com/pinbo/CRISgo) as published before [33]. For the restriction enzyme screen, we used A-genome specific primer pair CAPS-*WAPO-A1*-F and R1 and B-genome specific primer pair CAPS-*WAPO-B1*-F and R2 (S1 Table) followed by digestion with restriction enzyme *Xcm*I. The same primers and restriction enzyme digestion were used as Cleavage Amplified Polymorphism Sequence (CAPS) markers for subsequent generations.

### Generation of *WAPO1* transgenic lines

To validate the role of *WAPO-A1* in the regulation of SNS, we generated transgenic plants expressing the *WAPO-A1* gene driven by its native promoter. The genomic region of *WAPO-A1*, which included 4.8 kb upstream of the start codon, the complete coding region with its intron, and 1.5 kb downstream of the stop codon, was cloned into the hygromycin-resistance binary vector pLC41. Three different *WAPO-A1* alleles, each encoding a different WAPO-A1 protein (Table 1), were cloned and transformed into Kronos. The first genomic region of *WAPO-A*^*m*^*1* was obtained from the BAC library of diploid *T. monococcum* ssp. *monococcum* accession DV92, henceforth TmDV92 (genome A^m^) [34]. The WAPO-A^m^1 protein encoded by the TmDV92 allele differs from WAPO-A1 in polyploid wheat by three amino acids, but is otherwise similar to the H3 haplotype (Table 1). The second *WAPO-A1* genomic region was cloned from the BAC library of *T. turgidum* ssp. *durum* variety Langdon [35]. This construct, designated hereafter as LDN-C47, encodes the ancestral *Wapo-A1d* allele (H3), which has no deletion in the promoter region and the ancestral amino acids C47 and D384 (Table 1)

The third construct, named LDN-F47, was created by site-directed mutagenesis of construct LDN-C47 using primers described in S1 Table. The change from the ancestral cysteine (C47) into a phenylalanine (F47), resulted in a protein identical to the one encoded by the *Wapo-A1b* allele in the H2 haplotype (Table 1). However, the LDN-F47 clone differs from the natural H2 genomic region by three SNPs in the promoter region and two in the first intron [10]. Table 1 presents a summary of the different *WAPO-A1* constructs and their comparison with the endogenous *Wapo-A1a* allele (H1) present in the transformed variety Kronos.

To test the effect of the constitutive expression of *WAPO-A1*, we generated two additional *WAPO-A1* constructs using the same pLC41 vector. These constructs included the maize ubiquitin promoter (UBI), the synthesized *WAPO1* coding regions for the H3 (C47) and H2 (F47) haplotypes, and a MYC tag, hereafter referred to as UBI::C47:MYC and UBI::F47:MYC, respectively. The *WAPO1* coding region was synthesized by Genewiz into the pUC57 vector. The WAPO1 coding region was amplified using BP primers listed in Table S1. Amplified PCR product was gel extracted (Qiagen) and recombined into pDONRzeo using Life Technologies BP Clonase II following the manufacturer’s protocol. The pDONRzeo vector containing the desired WAPO1 coding region was recombined into the pLC41 vector using Life Technologies LR Clonase II, following the manufacturer’s protocol. Clones were verified by sanger sequencing at each cloning step.

Constructs were transformed into Kronos using *Agrobacterium-*mediated transformation (EHA105) at the UC Davis Plant Transformation Facility as described before [32]. Primers listed in S1 Table were used to genotype the transgenic plants and confirm the presence of the transgene. Transgenic T_0_ plants were advanced to T_1_ and characterized in the greenhouse for heading date, SNS, and spike and floral morphology. UBI::C47:MYC and UBI:F47 MIC transcript levels in leaves were determined by qRT-PCR using primers in Table S1. RNA extraction and expression analysis were done as previously described [36].

### Populations used to compare the effects of *WAPO-A1* haplotypes in the field

In our previous study, the *WAPO-A1b* H2 haplotype was associated with significantly higher SNS than the H1 and H3 haplotypes in common wheat, but the relative effect of H3 and H1 on SNS was not tested [10]. To compare the effect of the H1 and H3 haplotypes on SNS, we developed a population segregating for the H1 haplotype from Kronos and the H3 haplotype from the durum wheat variety Rusty (Table 1), which carried the same *Wapo-A1d* allele as Langdon [10]. From the Kronos x Rusty cross [37], 75 F_2_ plants were advanced to F_4_ by single seed descent (SDS) and genotyped for the *WAPO-A1* haplotype using a molecular marker previously developed for the *WAPO-A1* promoter deletion [10]. We selected eight heterozygous F_4_ plants (IDs= 12, 19, 51, 55, 69, 113, 120, 128) to generate eight Heterozygous Inbreed Families (HIFs). For each F_4:5_ HIF, we selected two homozygous sister lines (F_4:6_) – one fixed for H1 and the other fixed for H3.

The F_4:6_ grains were used for two field trials conducted at UC Davis experimental field station that were planted in October and harvested in June of the next year. The experiments will be referred using their harvest years in 2020 and 2021. Both field experiments were organized in a split plot, randomized complete block design (RCBD) with 10 replications, using the eight HIFs as main plots, and sister lines fixed for H1 and H3 haplotypes as subplots. We used single rows as experimental units in the 2020 experiment and small plots (3 rows = 1.86 m^2^) in the 2021 experiment. In 2021, we also measured heading date as the time from the first rain after sowing to the time when half of the plants in the plot have headed, grain number per spike (each replication was the average of 3 spikes), and total grain yield in kg/ha (small plots were harvested with a Wintersteiger Combine).

To compare the effect of the H2 relative to H1, we developed NILs segregating for these haplotypes in both tetraploid and hexaploid wheat. The tetraploid NILs were developed by introgressing the *WAPO-A1* H2 haplotype from UCD common wheat breeding line UC1110 (also referred to as CAP1, used as female) into tetraploid wheat variety Kronos using marker assisted backcrossing. After three backcrosses with Kronos, we self-pollinated the BC_3_ plants, and selected BC_3_F_2_ plants homozygous for the H1 and H2 haplotypes. The derived BC_3_F_3_ grains were sown at UC Davis in November 2020 in a completely randomized design (CRD) with nine replications, using 1-m rows as experimental units (10 spikes were measured per row and averaged). The hexaploid NILs were developed by backcrossing the H2 haplotype five times into the hexaploid line GID4314513 (H1) from the CIMMYT high-biomass program. We selected a high-biomass line as recurrent parent to increase the probability that increases in SNS and GNS would be translated into increases in grain yield. We developed BC_5_F_3_ homozygous NILs using GID4314513 as recurrent parent, and compared them in the field at UC Davis in 2021 using an RCBD with 10 replications and 3-row small plots (1.86 m^2^) as experimental units. We measured 4 spikes per plot (subsamples) and harvested the small plots with a combine.

### Expression studies

To test if the differences in SNS between H3 and H1 were associated with different levels of *WAPO-A1* expression, we compared the transcript levels of this gene in developing spikes of homozygous H1 and H3 sister lines derived from HIF #120. *WAPO-A1* transcript levels were determined by qRT-PCR using A-genome-specific primers UFO-A-RT-F2 and UFO-A-RT-R2 and PCR conditions described in our previous study [10]. Reactions were performed on an ABI 7500 Fast Real-Time PCR System (Applied Biosystems) using Fast SYBR GREEN Master Mix. Transcript levels were expressed as fold-*ACTIN* levels using the 2^ΔCT^ method.

Plants were grown in 1.4 L cones placed in CONVIRON growth chambers under 16 h light at 22 °C (330 mol intensity) and 8 h darkness at 17 °C for 25-30 days. For each replicate, we pooled 3-8 developing spikes from the main tiller when the lemma primordia were visible (W3.25) in the first experiment and when the floret primordia were present (W3.5) in the second experiment. Spike developmental stages are based on the Waddington scale [38]. We analyzed the two experiments separately and in a combined ANOVA using experiments as block.

To compare the transcript levels of class-B, -C and -E MADS-box genes in Kronos and the *wapo1* mutant, we extracted RNA from plants grown in a CONVIRON growth chamber under similar conditions as described above. RNA extraction and expression analysis were done as described previously [36]. Meristems were dissected when the plants were at the 6-7 leaf stage at the stamen primordia stage (∼W4.0). A total of 12 meristems were pooled per replicate.

### Statistical analyses

The interactions between *WAPO-A1* and *WAPO-B1* was tested using a 2 × 2 factorial ANOVA using homeologs as factors and alleles (WT and CRISPR mutants) as levels. The four possible homozygous classes – WT, *wapo-A1, wapo-B1*, and *wapo1* double mutant – were selected from the F_2_ progeny of an F_1_ plant from the cross *wapo1* x Kronos WT without the CRISPR-Cas9 transgene.

For the transgenic lines TmDV92, LDN-C47, and LDN-F47, we received three to five different transgenic events. Since the number of T_1_ non-transgenic sister lines was small for each event, we pooled the non-transgenic sister lines from the different events generated with the same construct. We then compared the different transgenic events with the pooled non-transgenic lines with the same construct using Dunnett tests. All statistical analyses were performed with SAS version 9.4. Homogeneity of variance was tested using the Levene’s test and normality of residual using the Shapiro-Wilks test as implemented in SAS v9.4. If necessary, data was transformed to restore the assumptions of the ANOVA.

## ACKNOWLEDGEMENTS

This project was supported by the Agriculture and Food Research Initiative Competitive Grants 2017-67007-25939 (WheatCAP) from the USDA National Institute of Food and Agriculture and by the Howard Hughes Medical Institute. DW is a Howard Hughes Medical Institute Fellow of the Life Sciences Research Foundation (http://www.lsrf.org/). We thank Mariana Padilla and Oswaldo Chicaiza for excellent technical assistance in designing, managing, and collecting all field experiment data and to Juan Debernardi for reviewing the manuscript and valuable suggestions. We also thank André Schönhofen, Xiaoqin Zhang, and Priscilla Glenn for their help with the introgression of the H2 haplotype from common wheat into Kronos and the hexaploid line GID4314513.

## AUTHOR CONTRIBUTION STATEMENT

SK and HL conducted most of the experimental work. SK wrote the first version of the manuscript. SC and DW contributed transgenic constructs with DW also contributing to experimental work and guidance to SK. JZ developed some of the near isogenic lines used in the study. CL contributed to the experimental work and supervised HL. JD initiated and coordinated the project, contributed to data analyses, and supervised SK. All authors reviewed the manuscript and provided suggestions.

## Notes

### Competing Interest Statement

The authors have declared no competing interest.

